# Ethanol-induced activation of BMP signaling and reprogramming of cardiomyocytes transcriptome

**DOI:** 10.64898/2026.07.16.738749

**Authors:** Jyoti Maddhesiya, Aditya Gautam, Hamim Zafar, Bhagyalaxmi Mohapatra

## Abstract

Congenital heart disease (CHD) comprises a diverse group of structural heart defects present at birth due to complex interactions between genetic and environmental factors. Prenatal alcohol exposure (PAE) is a known environmental factor that disrupts fetal cardiogenesis and increases the risk of CHD. However, the molecular mechanisms behind ethanol (EtOH)-induced CHD remain obscure. This study investigated the effects of EtOH on bone morphogenetic protein (BMP) signaling and transcriptomic reprograming in HL-1 cardiomyocytes. HL-1 cells were treated with varying concentrations of EtOH (25, 50, and 100 mM) for 24 h. 100 mM of EtOH exposure significantly enhanced SMAD1/5 phosphorylation and upregulated BMP-responsive genes, namely *Id1, Gata4, Mef2c,* and *Nkx2.5*. Increased histone acetyltransferase activity further validated activation of BMP signaling through histone hyperacetylation. These effects were reversed by the BMP pathway inhibitor LDN-193189, confirming pathway-specific activation. Further, transcriptome analysis following 100 mM EtOH treatment identified 3,876 differentially expressed genes. KEGG enrichment analysis revealed significant dysregulation of cardiogenic pathways, including TGF-β, Hedgehog, PI3K-Akt, Notch, FoxO, and calcium signaling pathways, along with extracellular matrix-receptor interaction and focal adhesion pathways. Gene Ontology analysis highlighted disturbances in heart development, cellular differentiation, apoptosis, extracellular matrix (ECM) organization, and chromatin regulation. Network analysis identified key hub genes, viz. *Kras, Fn1, Col1a1, Prkaca, Fbn1, Col6a1, Col6a2, Ccnd1, Col1a2* and *Myc* which are upregulated and *Hsp90aa1*, *Mdm2*, *Jun*, *Hras*, *Il6, Hsp90ab1, Pdgfra, Cdkn1a, Pparg, Fos and Hspa8* are downregulated which were subsequently validated by qRT-PCR. Collectively, these findings provide novel insights into the molecular basis of EtOH-induced CHD and identify potential biomolecule candidates for future therapeutic investigation.

## 1. Introduction

Congenital heart disease (CHD), the most common congenital anomaly worldwide, arises from structural abnormalities that develop during early embryonic cardiogenesis and remains a leading cause of infant morbidity and mortality. Cardiac development is a highly coordinated and intricately regulated process of embryogenesis that requires precise cellular migration, proliferation, differentiation, and morphogenetic remodeling. Multiple evolutionarily conserved signaling pathways, including TGF-β, BMP, WNT, NODAL, and FGF function in a tightly synchronized manner to ensure proper cardiac specification, patterning, and morphogenesis. Among the signaling pathways governing cardiogenesis, BMP signaling plays a pivotal role in cardiac lineage specification, heart tube formation, chamber morphogenesis, and myocardial differentiation (Garside *et al*., 2013). BMP ligands activate serine/threonine kinase receptors, leading to phosphorylation of SMAD1/5/8 and subsequent transcriptional activation of cardiac developmental regulators, including *Id1*, *Id2*, *Id3*, *Gata4*, *Nkx2-5*, *Irx4*, *Tbx5*, and *Mef2c* (Izumi *et al*., 2006; Weiskirchen *et al*., 2009; Du *et al*., 2010; Kantzer *et al*., 2022; Miyazono *et al*., 2005; Du *et al*., 2022; Cai *et al*., 2013; Chhabra *et al*., 2019). Any perturbation in the signaling networks during embryogenesis can disrupt cardiac morphogenesis and predispose to CHD. Although genetic defects account for a proportion of CHD cases, accumulating evidence indicates that complex interactions between genetic susceptibility and environmental exposures contribute to the majority of disease burden (Blue *et al*., 2012; Maddhesiya and Mohapatra, 2024). Nevertheless, the precise pathogenic mechanisms underlying the majority of CHD cases remain incompletely understood.

Alcohol is one of the well-known environmental threats reported to induce CHD (Liu *et al*., 2013). A case-control study demonstrated that 33-100% of fetal alcohol syndrome (FAS) probands exhibited CHD, with atrial septal defects (ASD) and ventricular septal defects (VSD) being the most frequently observed cardiac anomalies (Burd *et al*., 2007). Consistently, another case-control study conducted in Spain evaluating different patterns of maternal alcohol consumption and congenital anomalies reported a significantly increased risk of CHD exclusively among mothers who consumed the highest daily amounts of alcohol (≥92 g/day) (Martínez-Frías *et al*., 2004). Epidemiological evidence further indicates that alcohol intake during the periconceptional period is significantly associated with cardiac malformations, particularly transposition of the great arteries and outflow tract defects (Grewal *et al*., 2008). In a California-based cohort study, maternal alcohol consumption of less than once per week was associated with a 1.6 to 2.1-fold increased risk of d-transposition of the great arteries along with suggestive associations with neural tube defects (NTDs) and cleft lip with or without cleft palate (CLP) (Grewal *et al*., 2008). More recently, a case-control study reported that maternal alcohol consumption during the periconceptional period was associated with a threefold increased risk of CHD in offspring (Mateja *et al*., 2012). A comprehensive meta-analysis encompassing 55 studies reported that alcohol consumption by both mothers and fathers was significantly associated with an increased risk of CHD in offspring (Zhang *et al*., 2020). This analysis further demonstrated a statistically significant association between maternal alcohol exposure and tetralogy of Fallot (ToF).

Moreover, Animal model studies provide compelling evidence supporting the association between alcohol exposure and cardiac defects. A seminal mouse study revealed that alcohol exposure with high levels during gestational days 8, 9, and 10 induced ventricular septal defects (VSDs) in nearly 75% of embryos (WEBSTER *et al*., 1984). Consistently, another mouse study unveiled the association of prenatal alcohol exposure (PAE) with cardiac defects with an incidence of approximately 87% (Serrano *et al*., 2010). More recently, embryonic alcohol exposure (EAE) results in development of cardiomyopathy and diastolic dysfunction in adult zebrafish. Additionally, the RNA-sequencing analysis of EAE ventricles revealed altered expression of both novel genes and genes associated with heart failure, indicating long-term molecular consequences of early alcohol exposure (Weeks *et al*., 2024).

Human stem cell-based models have further strengthened these observations. A collaborative study on human-induced pluripotent stem cells (hiPSCs) have shown that alcohol exposure impairs cardiomyocyte differentiation in a dose-dependent manner and associated with mitochondrial dysfunction (Man *et al*., 2025). In line with this, Liu et al. (2021) through proteomic profiling demonstrated that chronic alcohol exposure in hiPSC-derived cardiomyocytes causes profound cellular and functional abnormalities, including increased cell death and oxidative stress, disrupted Ca²⁺ handling, abnormal action potentials, and reduced contractile function (Liu *et al*., 2021).

In this study, we investigated the impact of alcohol exposure on cardiomyocytes by treating cells with varying concentrations of alcohol, with a specific focus on disruption of BMP signaling. Furthermore, we employed transcriptome profiling to comprehensively delineate alcohol-induced molecular perturbations and to elucidate downstream gene regulatory networks involved in cardiac development with the aid of EtOH-associated biomolecules discovery.

## 2. Materials and Methods

### 2.1 Cell Culture and EtOH Treatment

The effect of EtOH was checked in HL-1 cardiomyocyte cells (purchased from Sigma-Aldrich, Cat No. SCC065). Cells were cultured in Claycomb Medium (Sigma-Aldrich, Cat No. 51800C) supplemented with 100 μM norepinephrine, 10% fetal bovine serum (FBS) and 4 mM L-glutamine, and maintained at 37°C in a humidified atmosphere containing 5% CO₂. At 80-90% confluency, cells were routinely sub-cultured. For experimental treatments, cells were seeded in T25 culture flasks and treated at 70-80% confluency. High-purity grade ethanol (EtOH) (Merck, USA) was used for all experiments.

LDN-193189 (Sigma-Aldrich, USA), a well-recognized dorsomorphin derivative and inhibitor of BMP signaling (Cuny *et al*., 2008), was employed (for validation of BMP pathway dependence of alcohol). LDN-193189 was prepared as a 1 mM stock solution in DMSO and diluted in PBS for all experimental use. LDN-193189 treatment was given 30 min prior to EtOH exposure.

The cells were categorized into four groups: EtOH treatment, alcohol plus LDN-193189 treatment, LDN-193189 treatment and un-treatment. The treated culture flasks were sealed completely to prevent EtOH evaporation.

## 2.1 Cell viability Assay

MTT assay was performed to evaluates the cytotoxicity of EtOH and LDN-193189 on cell viability and mitochondrial activity. HL-1 cardiomyocytes were seeded in 96-well culture plates (2 × 10⁴ cells per well) and allowed to adhere for 24 h. Cells were then treated with an increasing concentration of EtOH (25, 50, 100, 150, 200, 250, and 300 mM). In independent set of experiments, the effects of EtOH were assessed at 6, 12, 24, and 36 h for evaluating time-dependency.

### Cytotoxicity due to LDN-193189 (BMP signaling inhibitor)

In parallel, the cytotoxic effect of the BMP signaling inhibitor, LDN-193189 was investigated in a concentration range of 2-1000 nM (2, 4, 8, 16, 50, 100, 150, 200, 250, 500, 750, and 1000 nM). Following 24 h of treatment, the culture medium was removed and 10 µL of MTT solution was added, then incubated at 37°C for 4 h. Subsequently, the MTT solution was replaced with 100 µL of dimethyl sulfoxide (DMSO) in each well to solubilize the formazan crystals. Plates were incubated at 37°C in the dark for an additional 15 min. The formation of a purple/blue formazan product, indicative of mitochondrial dehydrogenase activity was quantified by measuring absorbance at 570 nm using a Synergy HTX multi-mode plate reader (BioTek Instruments, Inc., USA) (Riss *et al*., 2016). The half-maximal inhibitory concentration (IC₅₀) was estimated from dose-response curves, and percentage cell viability was calculated relative to untreated controls. All experiments were performed in three independent biological replicates, with technical triplicates for each condition. %Cytotoxicity = [ODcontrol - ODtreated*/*ODcontrol] ×100 % Cell viability = [100 - %Cytotoxicity]

## 2.2 Western blotting

To determine the appropriate EtOH concentration for subsequent experiments, the level of SMAD1/5 phosphorylation was assessed by Western blotting in HL-1 cardiomyocyte cells. Cells were seeded in T25 culture flasks and, after 24 h, treated with varying concentrations of EtOH (25 mM, 50 mM, and 100 mM). In a separate set of experiments, cells were co-treated with 100 mM EtOH and increasing concentrations of the BMP signaling inhibitor LDN-193189 (2, 4, 8, 16, 50, 100, 150, 200, 250, 500, 750, and 1000 nM). Following 24 h of EtOH and/or LDN-193189 treatment, whole-cell lysates were prepared using RIPA buffer and protein concentrations were quantified using the Bradford assay. Equal amounts of protein (25 µg) were mixed with Laemmli’s reducing sample buffer, denatured at 95°C for 5 min, and resolved on an 8% SDS-PAGE gel. Proteins were subsequently transferred onto a polyvinylidene difluoride (PVDF) membrane. Membranes were blocked with 5% non-fat dry milk in 1× TBST for 2 h at room temperature (RT), followed by incubation overnight at 4°C with a phospho-SMAD1/5 primary antibody (Cell Signaling Technology, USA). After washing three times with 1× TBST, membranes were incubated with HRP-conjugated goat anti-rabbit IgG secondary antibody (Genei, Germany) for 2 h at RT. Following five additional washes with 1× TBST (5 min each), immunoreactive bands were visualized using an enhanced chemiluminescence detection kit (GE Healthcare, USA) and imaged on an Amersham™ Imager 680 (Japan).

## 2.3 Real-Time PCR

To evaluate the effect of EtOH on BMP downstream signaling and cardiac-specific gene expression, real-time PCR (qRT-PCR) was performed. Total RNA was isolated from HL-1 cardiomyocytes using TRI Reagent (Merck, Germany), and RNA integrity was verified by 1% agarose gel electrophoresis. RNA concentration and purity were checked using a NanoDrop spectrophotometer (Thermo Fisher Scientific, USA). For cDNA synthesis, 2 µg of total RNA was treated with DNase I (Thermo Fisher Scientific, USA) and reverse transcribed using the RevertAid First Strand cDNA Synthesis Kit (Thermo Fisher Scientific, USA) with random hexamer primers. The expression of cardiac-enriched genes *Id1*, *Gata4, Nkx2.5 and Mef2c* was assessed at various concentrations (25 mM, 50 mM, and 100 mM) of EtOH exposure for 24 h using a QuantStudio™ 5 Real-Time PCR System (Applied Biosystems, CA, USA) and SYBR™ Green Master Mix (Sigma-Aldrich, USA). Based on time-course analysis, subsequent experiments were performed at the selected time point to assess the expression of cardiac transcription factors (*Gata4, Nkx2.5,* and *Mef2c*) and the BMP downstream target *Id1* following treatment with EtOH and/or LDN-193189. All qRT-PCR experiments were conducted in three independent biological replicates, each performed in technical triplicates. Gene expression levels were calculated using the comparative Ct (2⁻ΔΔCt) method, and results were expressed as fold change ± standard error of the mean (SEM).

## 2.4 HAT activity Assays

To assess the effect of EtOH on histone H3 acetylation in cardiomyocytes, histone acetyltransferase (HAT) activity was measured in HL-1 cells. Cells were seeded in T25 culture flasks and treated with 100 mM EtOH and/or 150 nM LDN-193189 for 24 h. Following treatment, nuclear extracts were prepared using sequential extraction with cytoplasmic extraction (CE) buffer (10 mM HEPES, 60 mM KCl, 1 mM EDTA, 0.075% NP-40, 1 mM DTT, and 1 mM PMSF) and nuclear extraction (NE) buffer (20 mM Tris-HCl, 420 mM NaCl, 1.5 mM MgCl₂, 0.2 mM EDTA, 25% glycerol, and 1 mM PMSF). Nuclear protein concentration was determined using the Bradford assay, and 25 µg of nuclear lysate was used for each reaction. HAT activity was quantified using the EpiQuik™ HAT Activity/Inhibition Assay Kit (Epigentek, USA) according to the manufacturer’s protocol. Absorbance was measured at 450 nm using a Synergy™ HTX multi-mode microplate reader (BioTek Instruments, Inc., USA). The results were expressed as relative HAT activity compared with untreated controls.

## 2.5 RNA Seq analysis

Total RNA was isolated from EtOH-treated (100 mM) and untreated HL-1 cardiomyocytes (n=3 per group) using TRI Reagent (Merck, Germany). RNA quantity and integrity were estimated using the Qubit™ 4 Fluorometer (Invitrogen™, USA), and all samples exhibited RNA integrity numbers (RIN) > 9.1. High-quality RNA samples were used for library preparation using NEB Directional RNA Library Prep kit followed by paired-end sequencing (2 × 75 bp) on an Illumina NextSeq 2000 platform. Reference transcriptome preparation for *Mus musculus* (GRCm38.p6) was performed using the rsem-prepare-reference function in RSEM (Li and Dewey, 2011). Read alignment and transcript quantification were carried out using STAR aligner (Dobin *et al*., 2013) integrated within the rsem-calculate-expression function. Normalization and differential gene expression analysis between EtOH-treated and control samples were performed using the DESeq2 (Love *et al*., 2014) package in *R*. Genes exhibiting an absolute expression change of >1.5-fold (log₂ fold change ≥ ±0.585) with an *adjusted p-value* (*padj*) <0.05 were defined as differentially expressed genes (DEGs).

## 2.6 Data analysis

Functional annotation and Gene Ontology (GO) enrichment analysis of DEGs were conducted using the ShinyGO web-based tool followed by visualization in *R*. Protein-protein interaction (PPI) network analysis was performed using the STRING database (Szklarczyk *et al*., 2019) with a high-confidence interaction score (≥0.9) and visualized in *R.* Hub genes were identified using the multiple topological properties of CytoHubba plugin (Chin *et al*., 2014) in Cytoscape v3.10.1 (Doncheva *et al*., 2019) including the Degree, Stress, Density of Maximum Neighborhood Component (DMNC), Maximum Neighborhood Component (MNC), Maximal Clique Centrality (MCC), Edge Percolated Component (EPC), Betweenness, Closeness, Radiality, Eccentricity, and Bottleneck.

## 2.7 RNA-Seq validation by qRT-PCR

The expression of top hub genes identified by RNA-seq was validated using qRT-PCR. RNA samples were isolated from independent EtOH-treated and untreated experiments. Complementary DNA (cDNA) synthesis and qRT-PCR reactions were performed in triplicate as described in the Materials and Methods (Section 2.4). For validation, five upregulated genes (*Kras, Fn1, Col1a1, Prkaca, and Ccnd1*) and five downregulated genes (*Hsp90aa1, Mdm2, Hras, Pdgfra, and Cdkn1a*) were selected. The expression level of genes was calculated using the comparative Ct (2⁻ΔΔCt) method, and results were plotted as fold change ± standard error of the mean (SEM). Statistical significance between EtOH-treated and untreated groups was evaluated using Student’s *t*-test, with *p*<0.05 considered statistically significant.

## 3. Results

### 3.1 Effect of EtOH and LDN-193189 on Cell viability

Cell viability analysis following EtOH exposure (25-300 mM) demonstrated a dose-dependent reduction in cell viability, with approximately 80% cell death observed after 250 mM EtOH treatment (Figure 1a). The IC₅₀ value for EtOH after 24 h of treatment was determined to be 175 mM. Furthermore, treatment with 100 mM EtOH resulted in a time-dependent decline in cell viability over a 6-36 h exposure period, with viability decreasing to approximately 50% at 36 h (Figure 1b). In contrast, LDN-193189 treatment (2-500 nM) exhibited a comparatively mild, dose-dependent cytotoxic effect, with more than 80% cell survival observed even at 1000 nM after 24 h (Figure 1c).

**Figure 1.**
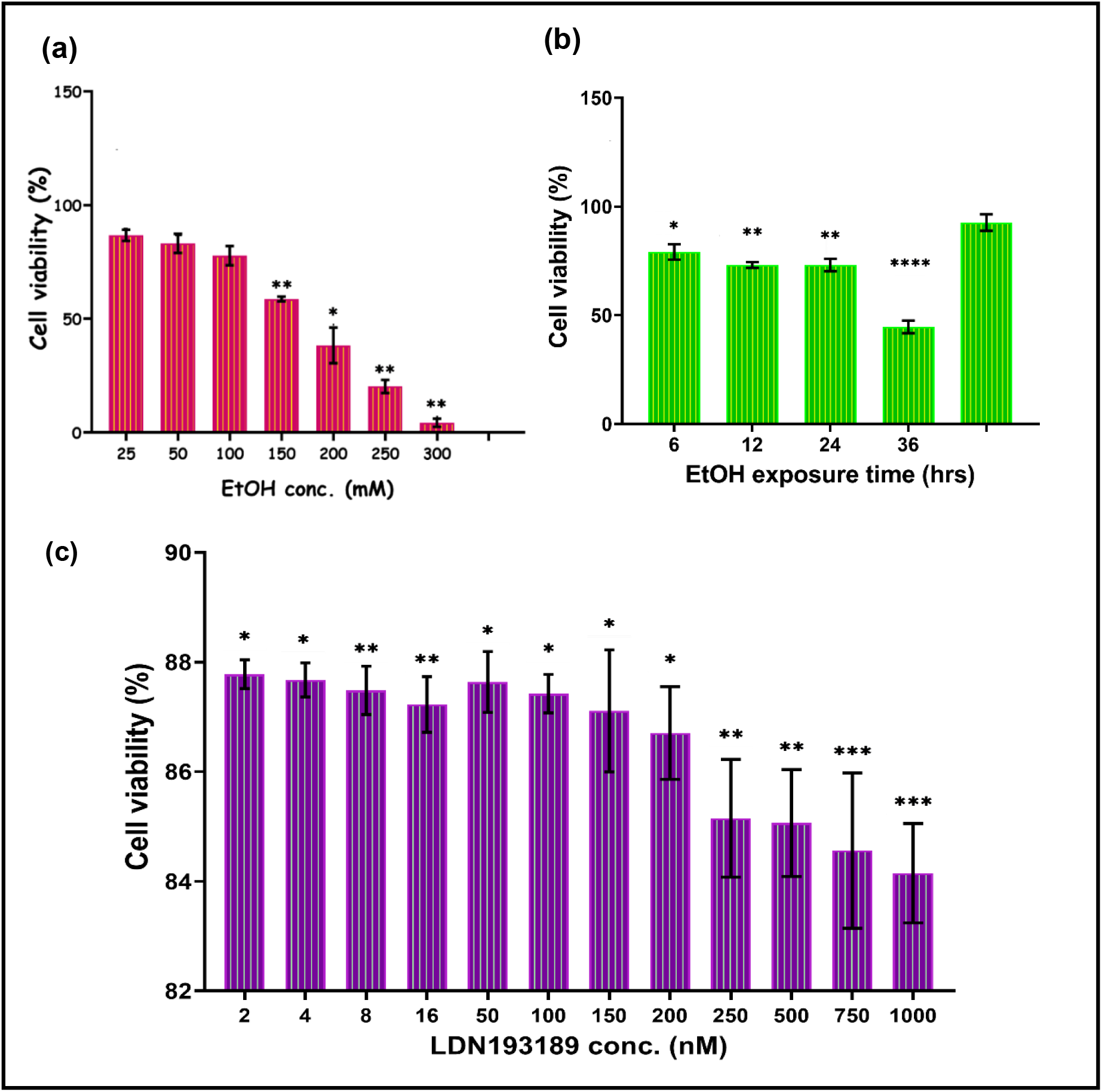
MTT Assay after EtOH **(a)** in dose-dependent manner **(b)** time-dependent manner and **(c)** LDN-193189 treatment in HL-1 cells: The effect of EtOH is measured in terms of cell viability in a dose (25-300 mM for EtOH, 2-1000 nM for LDN-193189) dependent manner for 24 h while 100 mM EtOH treatment (6-36 h). The error bar shows the mean and SD. The paired t-test was used to measure statistical significance

Morphological assessment using phase-contrast microscopy revealed that HL-1 cells displayed evident stress and altered morphology following 24 h of EtOH treatment, whereas untreated cells maintained normal cellular architecture. (Supplementary Figure S1).

### 3.1 Increased phosphorylation level of SMAD1/5 and inhibitory effect of LDN-193189 on EtOH induced SMAD1/5 phosphorylation

To evaluate the effect of EtOH on the BMP signaling pathway in HL-1 cells, phosphorylation of SMAD1/5 was checked by Western blotting. Cells were exposed to increasing concentrations of EtOH (25 mM, 50 mM, and 100 mM) for 24 h, which resulted in a dose-dependent increase in SMAD1/5 phosphorylation. Compared with untreated controls, SMAD1/5 phosphorylation was elevated by 4.29-fold at 25 mM (p=0.0666), 12.02-fold at 50 mM (p<0.0001), and 19.10-fold at 100 mM EtOH (p<0.0001) (Figure 2a-b). Based on this robust activation, 100 mM EtOH was selected for subsequent experiments.

**Figure 2.**
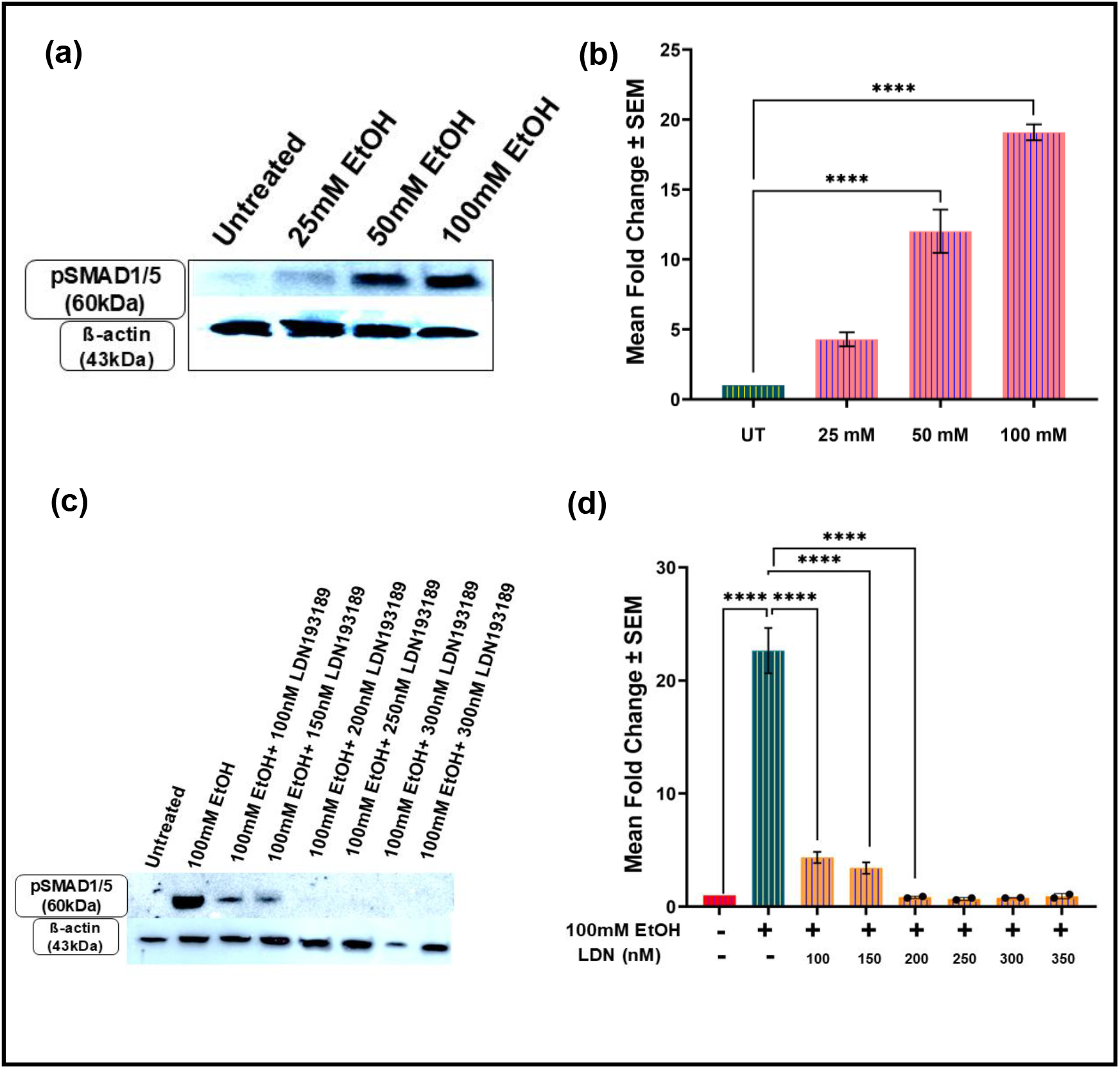
(a and. **b)** Effect of EtOH on the phosphorylation of SMAD1/5 in HL-1 cells. Western blotting illustrated the expression level of pSMAD1/5 was significantly enhanced after EtOH treatment. **(c and d)** EtOH-induced pSMAD1/5/8 phosphorylation was significantly prevented after LDN-193189 treatment. Statistical analysis-One-way ANOVA and *p<0. 05, **p<0.01, and ****p<0.0001.

To confirm the specificity of EtOH-mediated BMP pathway activation, HL-1 cells were co-treated with 100 mM EtOH and increasing concentrations of the BMP signaling inhibitor LDN-193189 (100-1000 nM). EtOH-induced SMAD1/5 phosphorylation was significantly attenuated in a dose-dependent manner following LDN-193189 treatment. A reduction of 5.20-fold at 100 nM, 6.61-fold at 150 nM, and a marked suppression (27.11-fold; p<0.0001) at concentrations ≥200 nM was observed compared to EtOH-treated cells (Figure 2c-d). Based on effective inhibition with minimal cytotoxicity, 150 nM LDN-193189 was selected for all subsequent experiments.

### 3.2 EtOH induced and LDN-193189 inhibitory effect on the expression level of cardiac specific transcription factor in dose-dependent manner

The effect of increasing concentrations of EtOH (25 mM, 50 mM, and 100 mM) on the expression of cardiac-enriched genes (*Id1, Gata4, Nkx2.5,* and *Mef2c*) was evaluated using qRT-PCR after 24 h of exposure. Overall, EtOH treatment resulted in a dose-dependent upregulation of all assessed genes. The expression of *Id1* was significantly increased by 1.64-fold (p=0.0055) and 2.59-fold (p<0.0001) following exposure to 50 mM and 100 mM EtOH, respectively, whereas a modest but non-significant increase (1.35-fold; p=0.1544) was observed at 25 mM EtOH (Figure 3a). Similarly, *Gata4* expression was significantly elevated across all EtOH concentrations, with increases of 1.60-fold (p=0.0002) at 25 mM, 1.94-fold (p<0.0001) at 50 mM, and 2.51-fold (p<0.0001) at 100 mM EtOH (Figure 3b). Notably, the master cardiac transcription factor *Nkx2.5* exhibited a robust and dose-dependent induction, with expression levels increased by 3.26-fold, 4.15-fold, and 5.69-fold following 25 mM, 50 mM, and 100 mM EtOH treatment, respectively (Figure 3c). Likewise, *Mef2c* expression was enhanced significantly by 2.60-fold (p<0.0001) at both 50 mM and 100 mM EtOH, whereas the increase at 25 mM EtOH (1.21-fold; p=0.7045) was not statistically significant (Figure 3d). To determine whether EtOH-mediated transcriptional activation occurs via BMP signaling, the effect of the BMP pathway inhibitor LDN-193189 (150 nM) was determined in 100 mM EtOH-treated HL-1 cells. Compared with untreated controls, 100 mM EtOH significantly increased the expression of *Id1*, *Gata4*, *Nkx2.5*, and *Mef2c* by 2.03-fold, 1.93-fold, 1.90-fold, and 1.70-fold, respectively (all p<0.0001) (Figure 3e-f). Importantly, co-treatment with LDN-193189 resulted in a significant reduction in EtOH-induced gene expression, with decreases of 1.67-fold for *Id1* (Figure 3e), 1.44-fold for *Gata4* (Figure 3f), 1.58-fold for *Nkx2.5* (Figure 3g), and 1.38-fold for *Mef2c* (all p<0.0001) (Figure 3h) relative to EtOH alone. Furthermore, when compared to untreated cells, the EtOH+LDN-193189 group still exhibited a modest but significant increase in the expression of *Id1* (1.29-fold; p=0.0020) (Figure 3e), *Gata4* (1.34-fold; p=0.0013) (Figure 3f), and *Mef2c* (1.23-fold; p=0.0011) (Figure 3h). In contrast, the increase in *Nkx2.5* expression (1.20-fold; p=0.0770) (Figure 3g) did not reach statistical significance.

**Figure 3.**
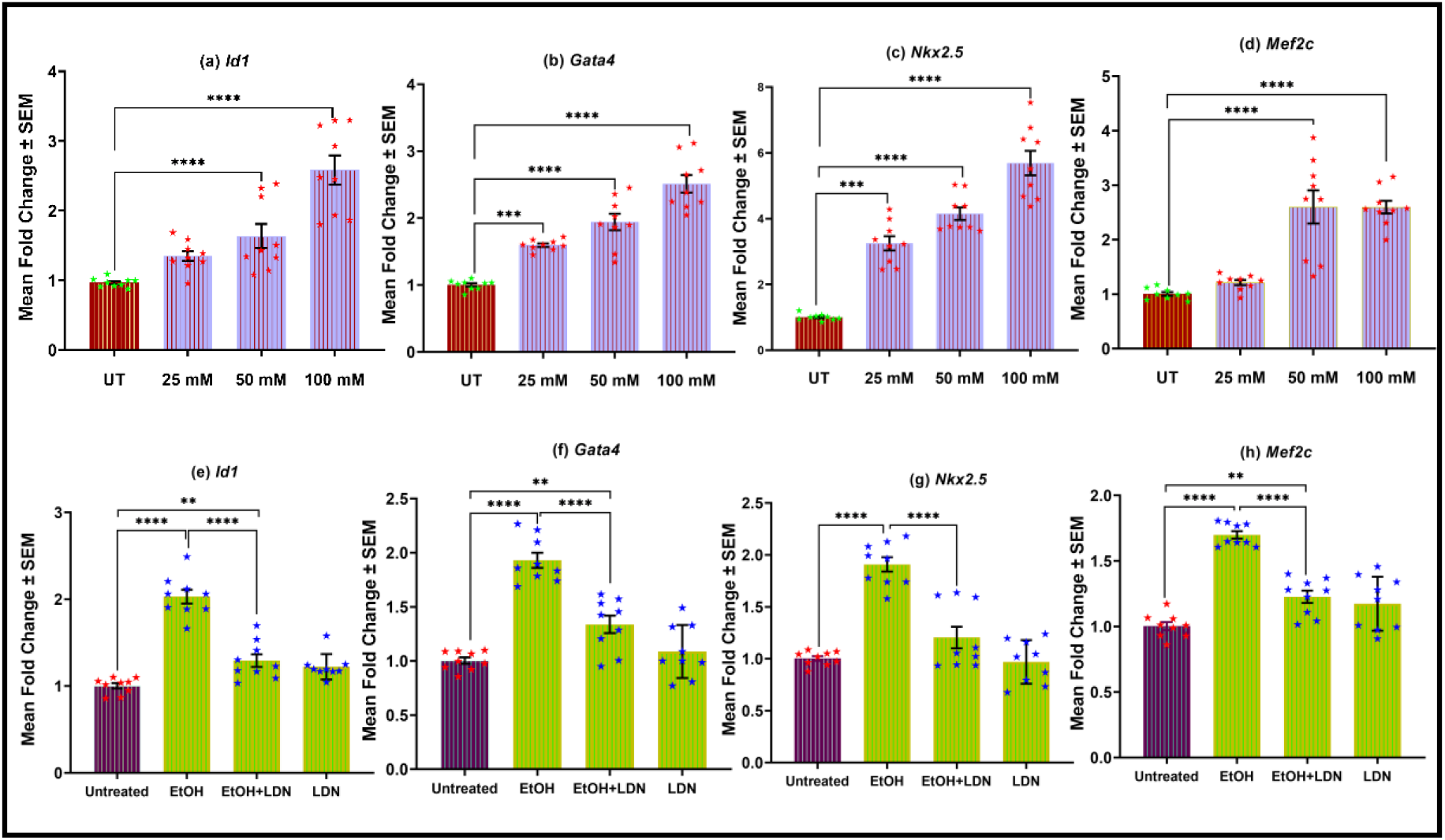
Effect of various (25, 50 and 100 mM) concentration of EtOH on the expression level of cardiac-specific genes. Real-time PCR showed that the expression of **(a)** Id1, **(b)** Gata4, **(c)** Nkx2.5 and **(d)** Mef2c was increased significantly after EtOH treatment in HL1-cells. Reduced expression of **(e)** Id1, **(f)** Gata4, **(g)** Nkx2.5 and **(h)** Mef2c after LDN-193189 treatment. Unpaired t-test and **p <0.01

### 3.3 Enhanced activities of HAT in HL-1 cells after EtOH exposure

Histone H3 acetylation in cardiomyocytes following EtOH and LDN-193189 exposure was evaluated using a histone acetyltransferase (HAT) activity assay after 24 h of treatment. A significant 2.26-fold increase in HAT activity (p=0.0029) (Figure 4a) was observed in HL-1 cells treated with 100 mM EtOH compared with untreated controls. Notably, HAT activity was significantly reduced by 1.76-fold (p=0.0066) (Figure 4a) when EtOH-induced HL-1 cells were co-treated with 150 nM LDN-193189. In contrast, treatment with LDN-193189 alone did not result in any significant change in HAT activity relative to untreated cells (Figure 4a).

**Figure 4.**
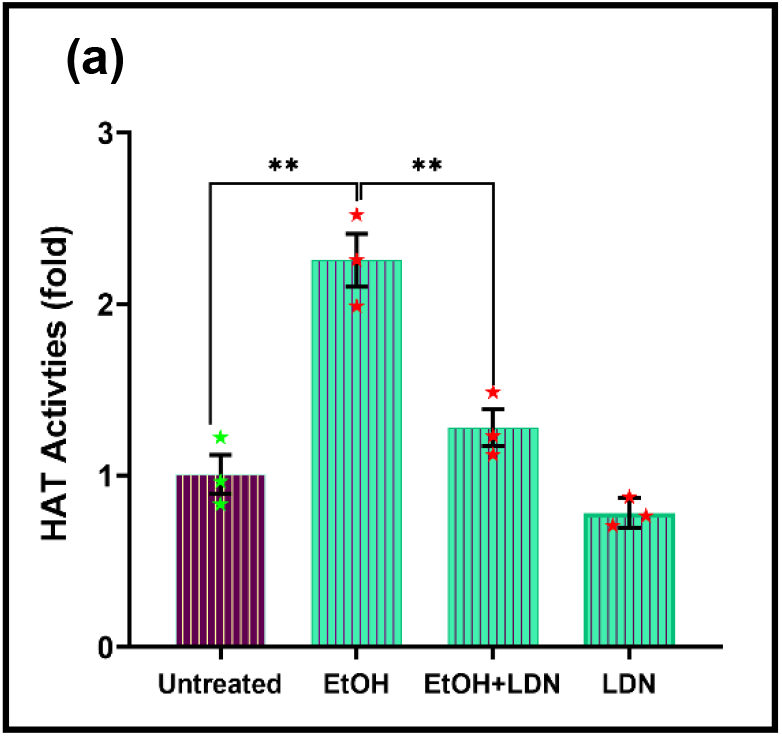
(a) Effects of EtOH and LDN-193189 treatment on HAT/HDAC activities in HL-1 cells. The HAT activities are increased significantly after alcohol exposure and reversed after the EtOH and LDN-193189 exposure. Unpaired t-test and **p <0.01

### 3.4 Alteration in the transcriptome profile of HL-1 cells after EtOH treatment

#### 3.4.1 Differentially expressed genes (DEGs) in EtOH-treated group

To further investigate the impact of EtOH exposure on HL-1 cardiomyocytes, bulk RNA sequencing (RNA-Seq) was performed to compare EtOH-treated cells (100 mM for 24 h) with untreated controls (*n*=3 per group). A total of 42,396 annotated genes were quantified based on the mouse genome (*Mus musculus*, GRCm38.p6) reference transcriptome. Among these, 15,919 genes were identified as differentially expressed between the EtOH-treated and untreated groups. Using a stringent threshold of *padj* <0.05 and ≥1.5-fold change, we identified 3,876 differentially expressed genes (DEGs). Of these, 1,717 genes (44.30%) were significantly upregulated, while 2,159 genes (55.70%) were significantly downregulated (Table 1).

**Table 1.** List of DEGs identified in between EtOH-treated and untreated group.

| List | No. of genes |
| --- | --- |
| Total DEGs | 15919 |
| Total DEGs (adjusted p-value <0.05) | 3876 |
| Total DEGs ($\geq 1.5$ fold change) | 3876 |
| Upregulated | 1717 |
| Downregulated | 2159 |

Principal component analysis (PCA) revealed a clear separation between EtOH-treated and untreated samples, highlighting robust transcriptional alteration following EtOH exposure (Figure 5a). A volcano plot illustrated the distribution of DEGs on the basis of fold change and statistical significance (Figure 5b). Additionally, a heatmap further demonstrated distinct expression patterns of DEGs across the samples of both groups (Figure 5c). To assess the clinical relevance of the identified DEGs, we intersected the EtOH-responsive genes with a curated CHD gene panel. This analysis revealed an overlap of 295 upregulated genes between the CHD panel and EtOH-induced upregulated genes, as well as 245 downregulated genes overlapping with the EtOH-induced downregulated genes and the CHD panel (Figure 5d).

**Figure 5.**
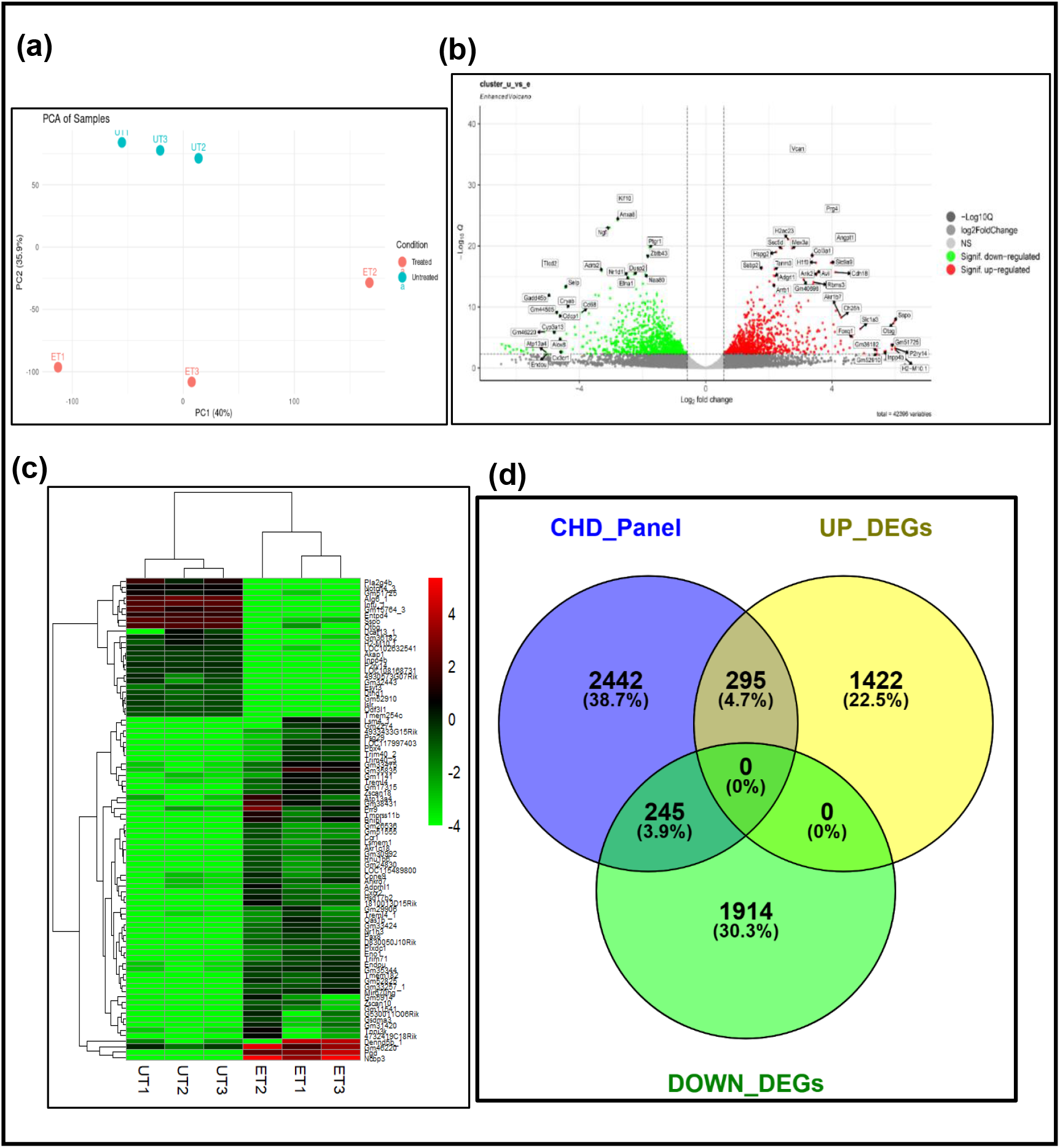
(a) This plot illustrates the variance in gene expression between the EtOH-treated and untreated group. The x-axis (PC1) and y-axis (PC2) show the principal components that explain the highest variance. EtOH-treated group are represented by red circles, and untreated group are represented by green circles, indicating distinct clustering between the two groups. **(b)** Volcano plot represents the DEGs between EtOH-treated and untreated group. x-axis represents the fold change and the y-axis represents statistical significance (-Log10 Q) for each gene. Red and green dots indicate significantly upregulated and downregulated genes respectively. **(c)** Heatmap illustrates the DEGs with >4 fold change between EtOH-treated and untreated group. The abscissa represents different samples: red color highlights upregulation; green color indicates downregulation. **(d)** Venn diagram demonstrates the intersection of upregulated (yellow circle) and downregulated (green circle) gene from our dataset with established cardiac developmental genes (blue circle), together with the corresponding list of shared genes

#### 3.4.1 KEGG pathway analysis for functional enrichment

The functional pathways affected by EtOH exposure was identified by KEGG pathway enrichment analysis for both upregulated and downregulated genes independently. The chord plot (Figure 6a) of upregulated genes revealed enrichment of several key signaling pathways of cardiogenesis, including TGF-β signaling, Hedgehog signaling, ECM-receptor interaction, AGE-RAGE signaling in diabetic complications, focal adhesion, Hippo signaling, Cellular senescence, PI3K-Akt signaling, Wnt signaling, Adrenergic signaling in cardiomyocytes, Tight junction, Calcium signaling, and MAPK signaling pathways.

**Figure 6.**
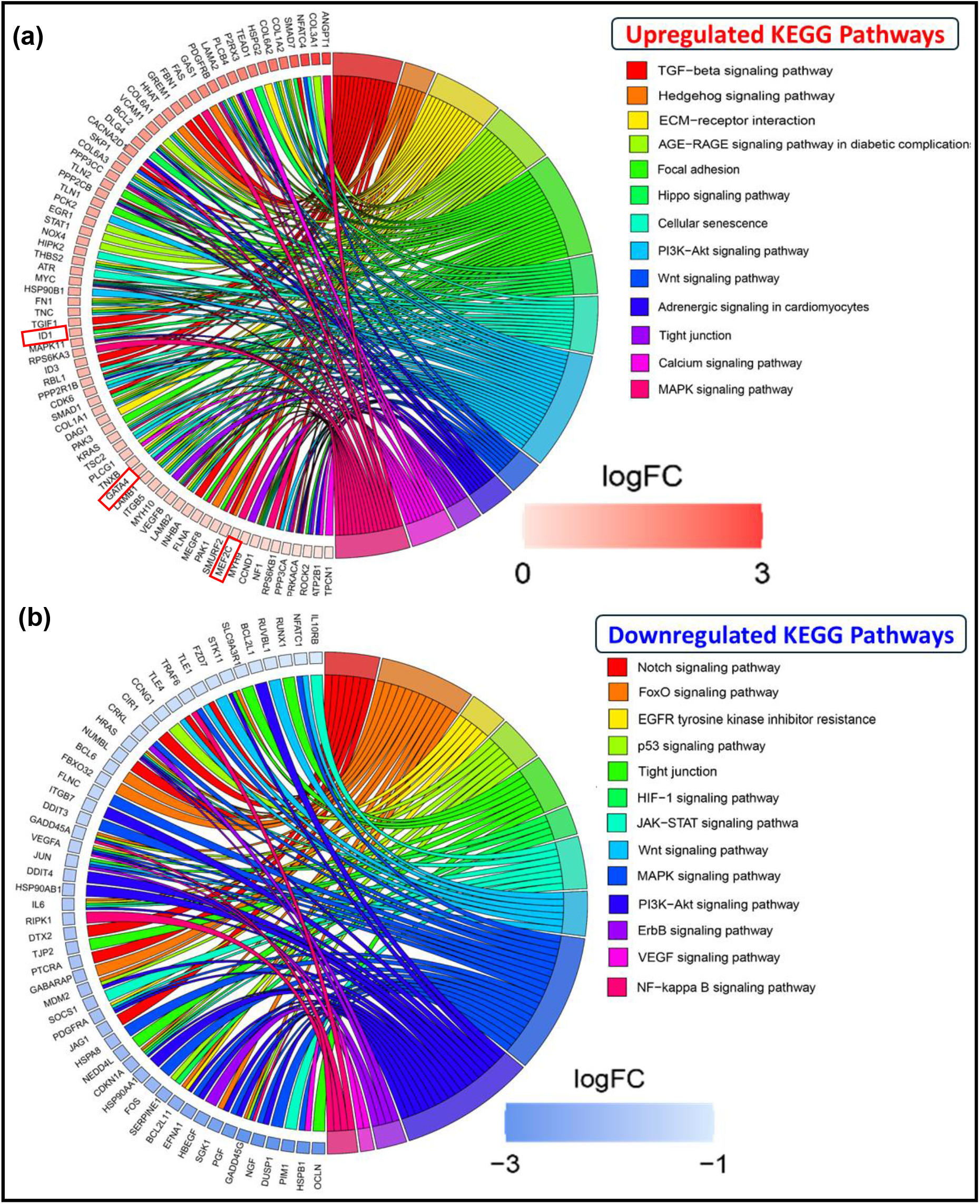
**KEGG Pathway enrichment analysis**: Chord plot illustrates enriched signaling pathways related to cardiogenesis **(a)** upregulated **(b)** downregulated pathways

Conversely, the chord plot of downregulated genes (Figure 6b) illustrated the suppression of multiple pathways crucially involved in cardiogenesis and cardiac homeostasis. These included the Notch signaling pathway, FoxO signaling pathway, EGFR tyrosine kinase inhibitor resistance, p53 signaling pathway, Tight junction, HIF-1 signaling pathway, JAK-STAT signaling pathway, Wnt signaling pathway, MAPK signaling pathway, PI3K-Akt signaling pathway, ErbB signaling pathway, VEGF signaling pathway, and NF-κB signaling pathway.

#### 3.4.2 Gene Ontology (GO) enrichment analysis

The effect of EtOH exposure on biological processes (BP), cellular components (CC), and molecular functions (MF) was determined by Gene Ontology (GO) enrichment analysis separately for upregulated and downregulated genes. The upregulated genes were enriched in 1,000 GO terms related to biological processes, 186 GO terms corresponding to cellular components, and 113 GO terms related to molecular functions. The top 20 significantly enriched GO terms of each category (BP, CC, and MF), ranked by fold enrichment, are illustrated in (Figure 7i-a-b-c).

**Figure 7.**
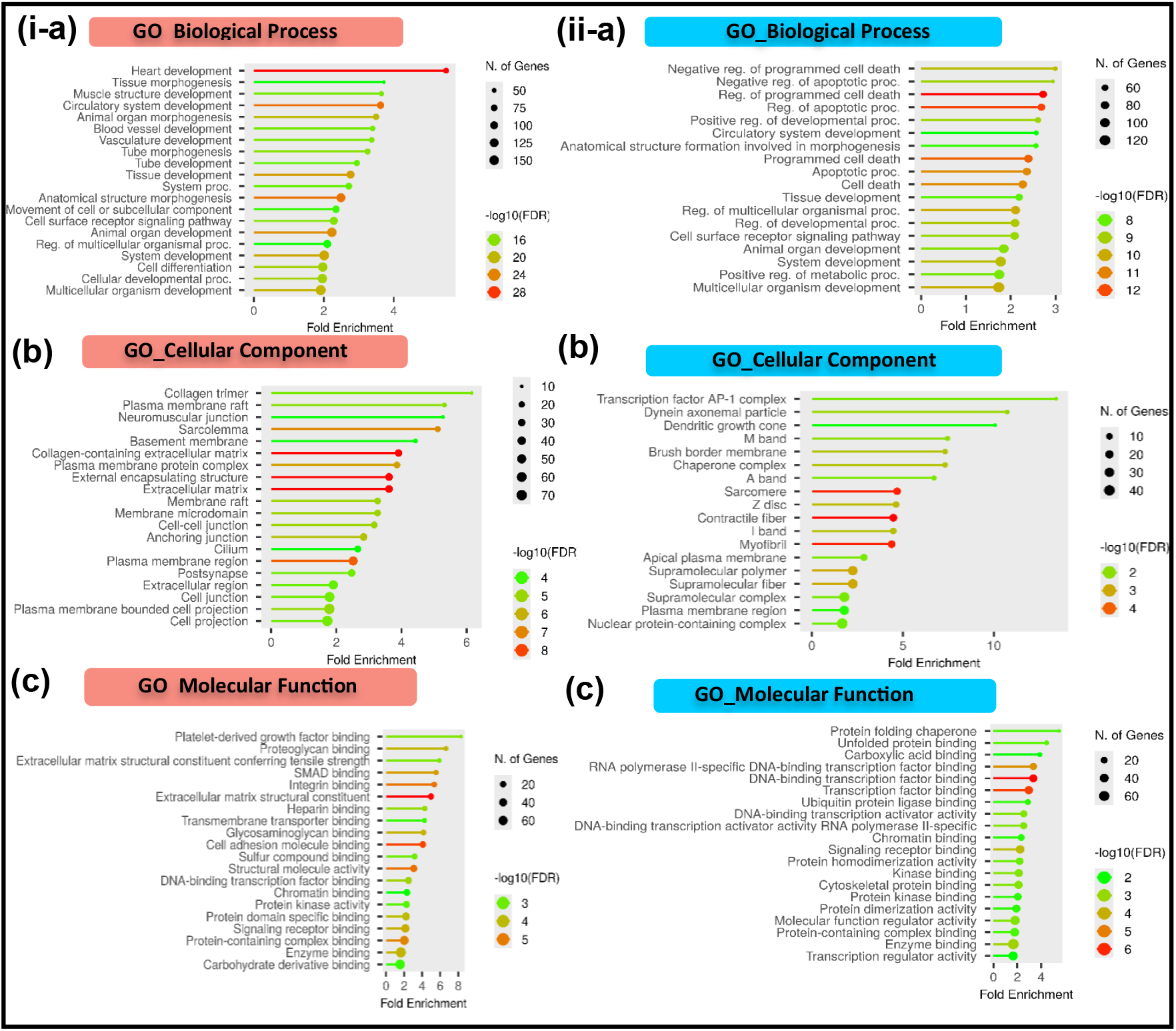
GO enrichment analysis. **(i)** upregulated **(ii)** downregulated terms **(a)** *Biological Processes: shows regulation of metabolic pathways crucial for cellular function and cardiac development* **(b)** *Cellular Component: reveals key roles in transcription factor binding and kinase activity* **(c)** *Molecular Function: demonstrates key roles in transcription factor binding and kinase activity*.

Similarly, downregulated genes are enriched in 1,000 GO terms under biological processes, 51 GO terms under cellular components, and 119 GO terms under molecular functions. The top 20 significantly enriched GO terms across BP, CC, and MF categories are presented in (Figure 7ii-a-b-c).

#### 3.4.3 Interactome analysis and hub gene identification

Additionally, a protein-protein interaction (PPI) network of cardiac-enriched upregulated genes (n=295) was constructed using the STRING database with a high confidence score of 0.700. The resulting network comprised 92 seed nodes, 818 interacting nodes, and 1,233 edges, representing interactions among the common differentially expressed genes (Figure 8a).

**Figure 8.**
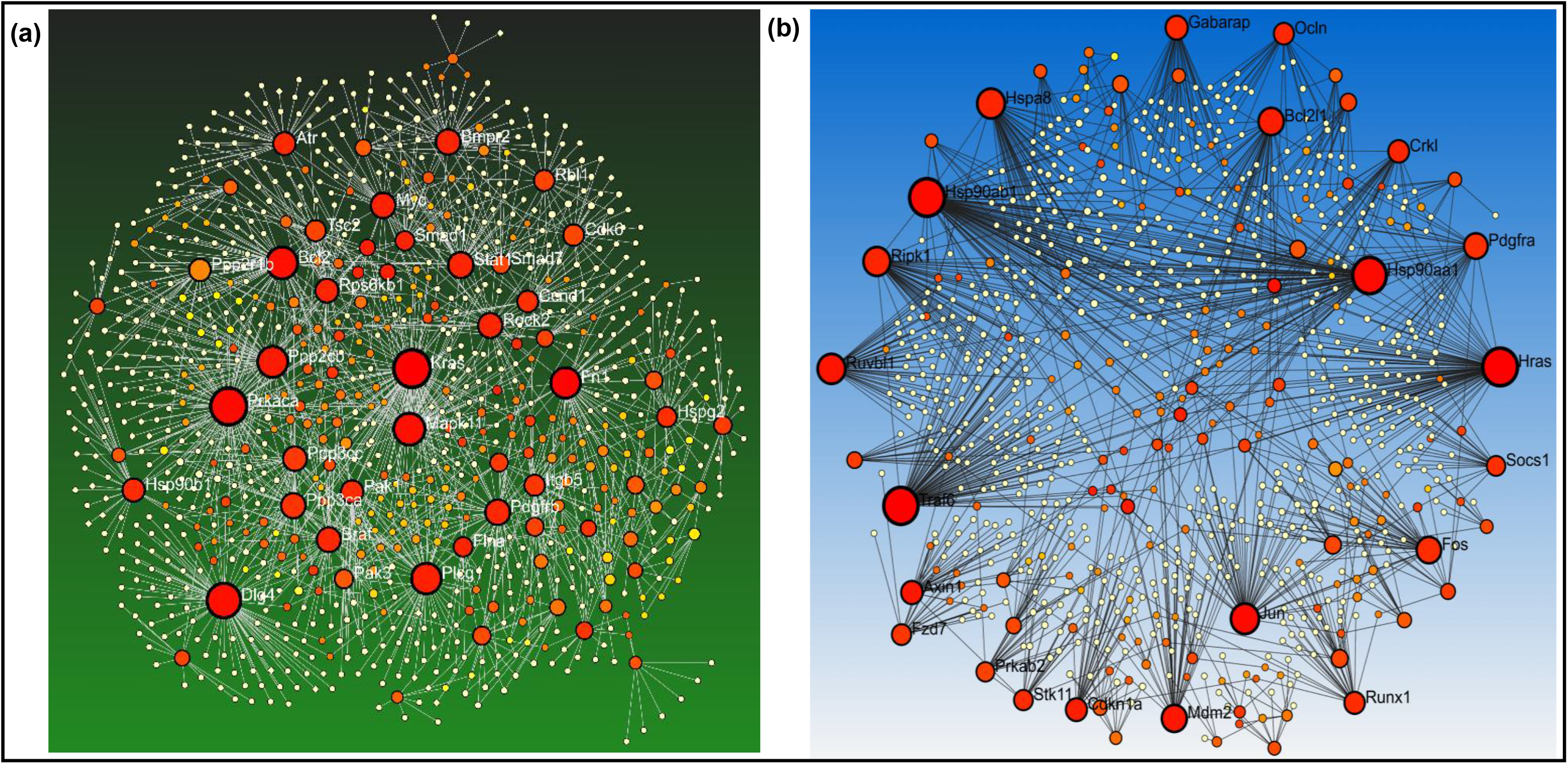
Protein-protein interactome: **(a)** upregulated **(b)** downregulated panels show the interacting partners of cardiac expressing genes using String database. The network nodes represent individual genes, and edges signify interactions between these genes. Size of node represents the degree of connectivity while gradient color of node indicates Betweenness

Similarly, a PPI network for cardiac-enriched downregulated genes (n=245) was generated using the same STRING confidence threshold, yielding a network of 121 seed nodes, 980 nodes, and 1,490 edges (Figure 8b).

To identify key regulatory genes within these networks, topological analysis was performed using the CytoHubba plugin in Cytoscape. Multiple algorithms-including Degree, Stress, Density of Maximum Neighborhood Component (DMNC), Maximum Neighborhood Component (MNC), Edge Percolated Component (EPC), Maximal Clique Centrality (MCC), Betweenness, Closeness, Radiality, Bottleneck, and Eccentricity-were applied. The results across these parameters were integrated using the Upset plot (Figure 9a-b) to identify the most consistently ranked nodes. The top 10 influential upregulated hub genes were identified (Table 2) based on this comprehensive analysis.

**Figure 9.**
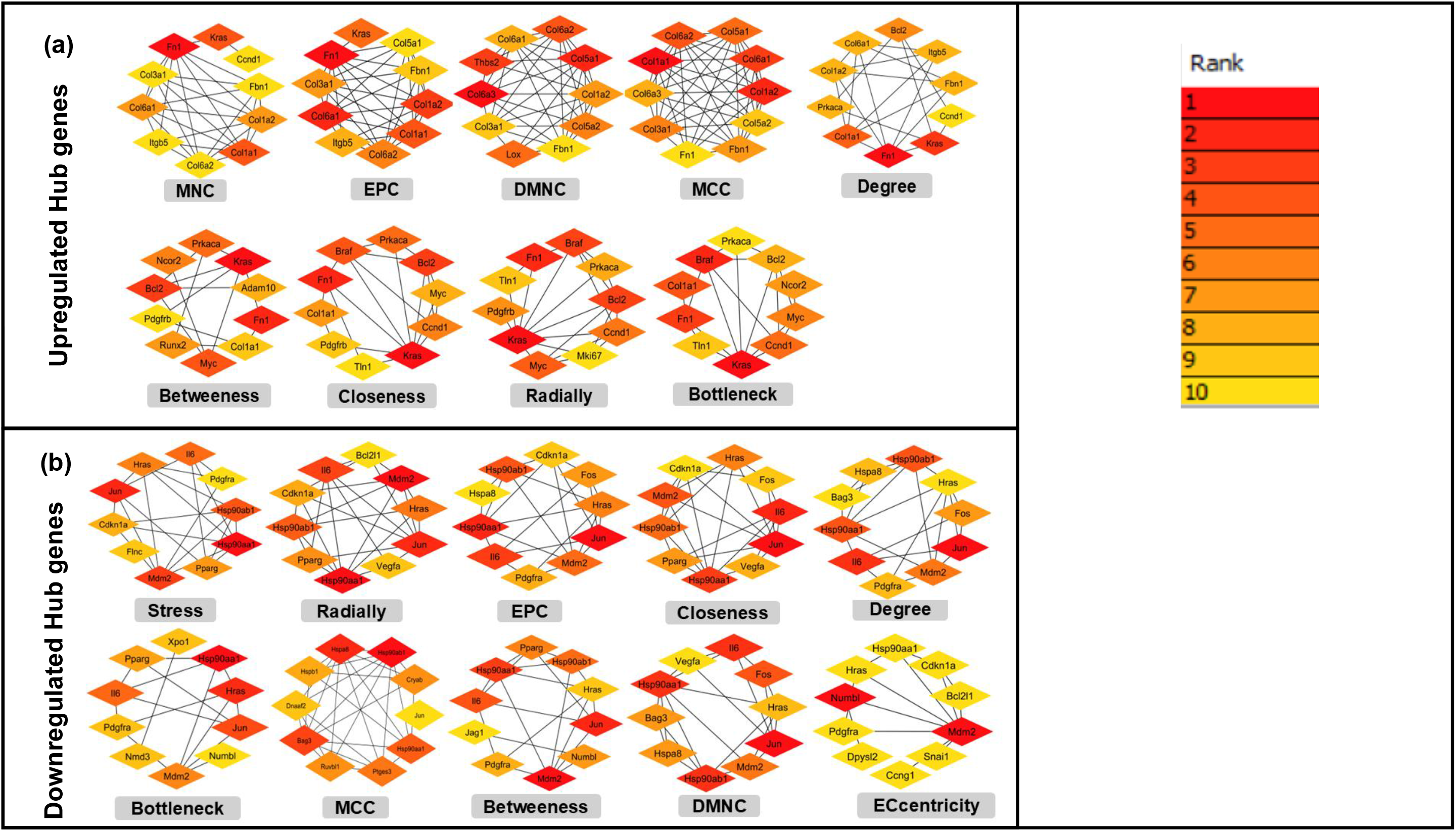
Upset plot showing the HubG’s networks of gene. **(a)** for upregulated **(b)** downregulated in different cytoHubba topological parameters including MNC, EPC, DMNC, MCC, Degree, Betweeness, Closeness, Radially and Bottleneck Stress, and ECcentricity. Darker nodes illustrating the higher rank of connectivity

**Table 2.**
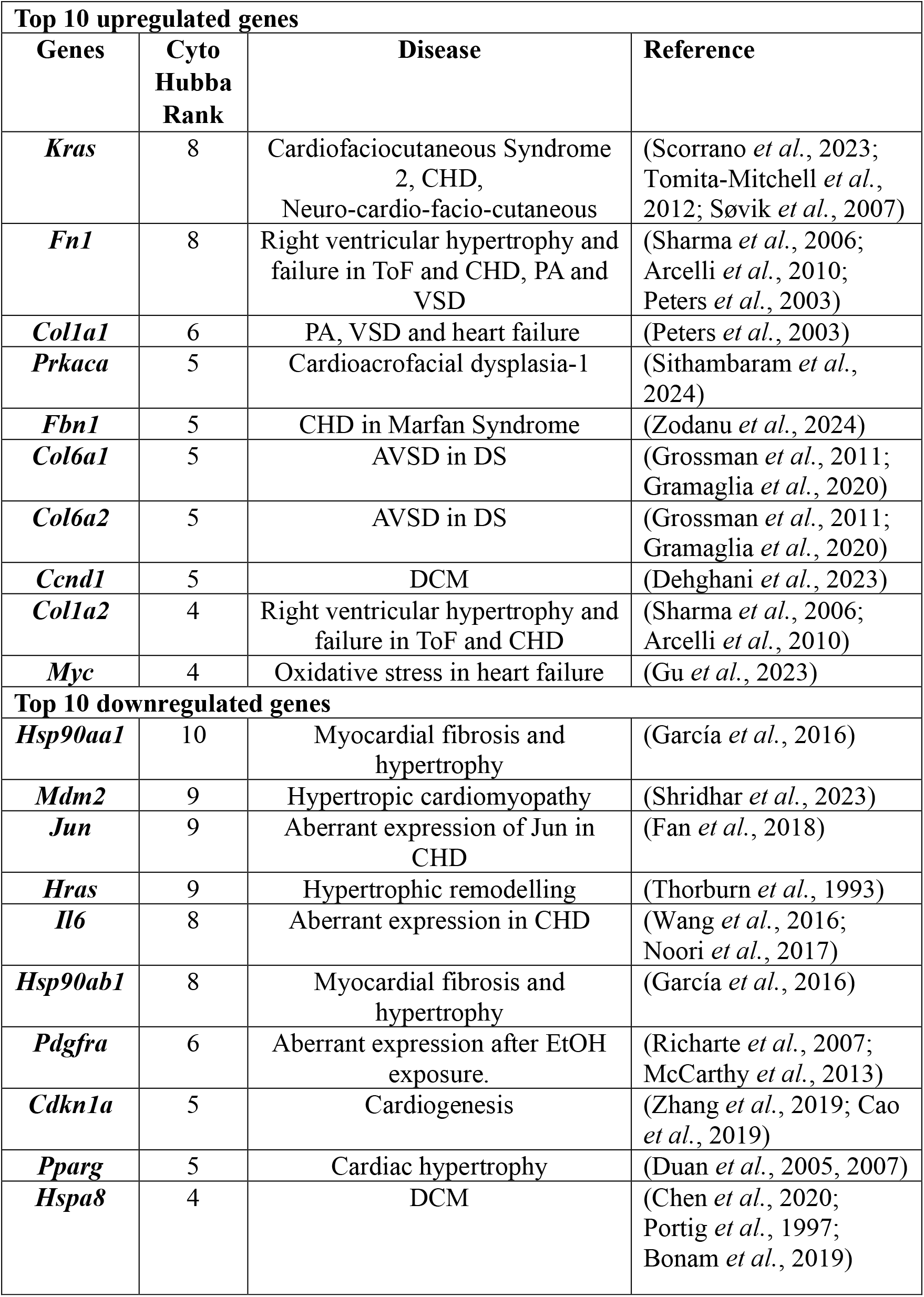
Represents the top 10 up-and down-regulated EtOH biomolecules for CHD.

Parallely, application of the same CytoHubba parameters to the downregulated PPI network result in the identification of the top 10 downregulated hub genes (Table 2), depicting key molecular players potentially involved in EtOH-mediated cardiac dysregulation.

### 3.5 Validation of top hub genes by qRT-PCR

qRT-PCR data analysis of top hub genes revealed an increase in the expression of *Kras* (3.67-fold, p=0.004), *Fn1* (3.57-fold, p=0.001), *Col1a1* (7.59-fold, p=0.002), *Prkaca* (4.21-fold, p=0.001), *Ccnd1* (1.61-fold, p=0.003), and decrease in the expression level of *Hsp90aa1* (6.76-fold, p=0.002), *Mdm2* (5.6-fold, p=0.001), *Hras* (3.81-fold, p=0.004), *Pdgfra* (2.38-fold, p=0.003), *Cdkn1a* (2.87-fold, p=0.001) in EtOH-treated group (Figure 10).

**Figure 10.**
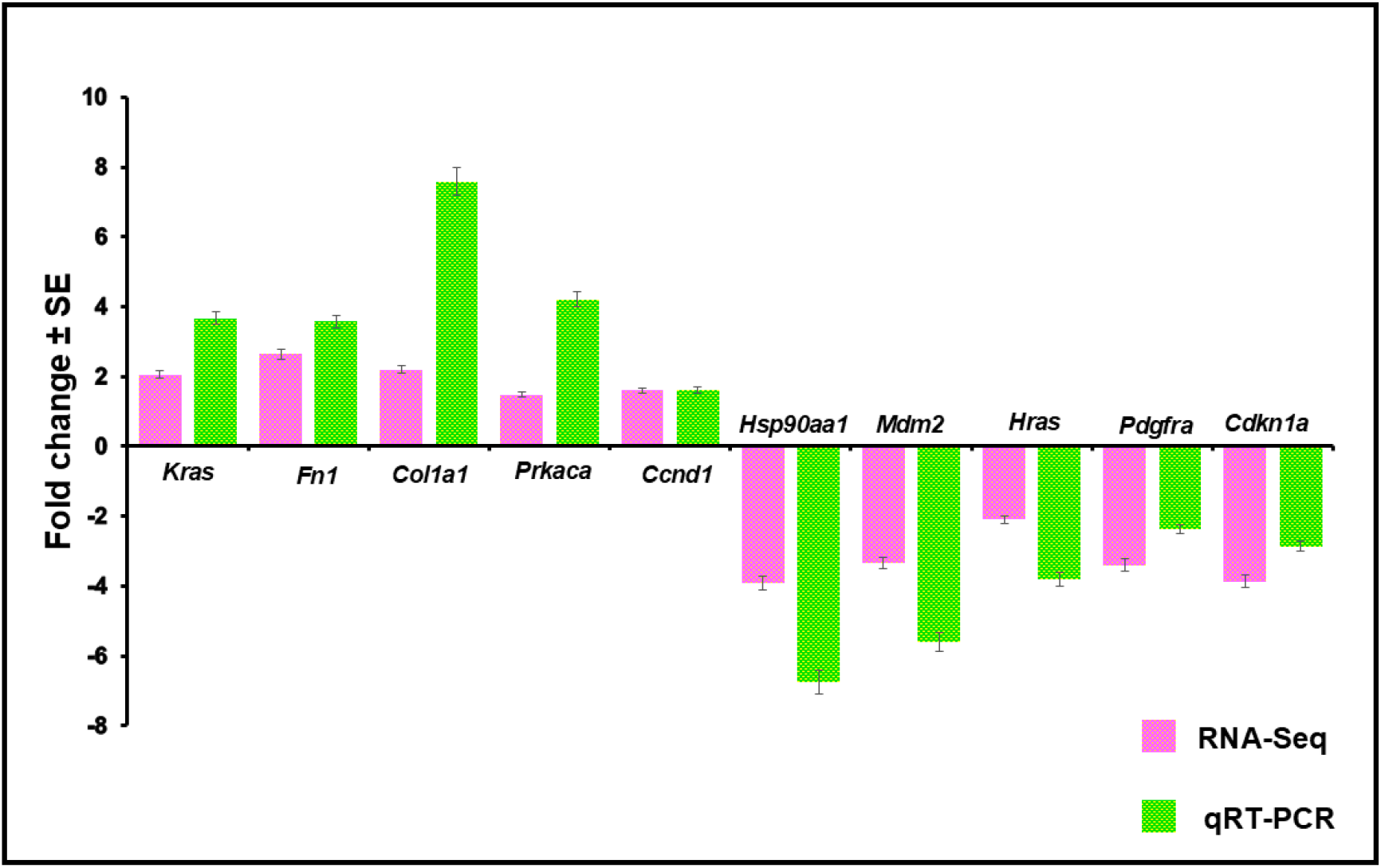
Bar diagram represents expression level of top5 upregulated genes **(Kras, Fn1, Col1a1, Prkaca, and Ccnd1)** and top5 downregulated genes **(Hsp90aa1, Mdm2, Hras, Pdgfra and Cdkn1a)**

## 4. Discussion

A large number of studies (both *in vivo* and *in vitro*) have demonstrated the abusive effects of alcohol, however the exact mechanism underlying alcohol-induced cardiotoxicity demands further investigation. Emerging evidence suggests that epigenetic alteration, particularly histone hyperacetylation, plays an important role in inducing alcohol-associated cardiac abnormalities. Several studies have revealed the alcohol-induced hyperacetylation of genes involved in cardiac development, leading to aberrant gene expression profiles (Zhang *et al*., 2014; Gao *et al*., 2015; Peng *et al*., 2015). Notably, Shi et al. (2016) showed that alcohol exposure in cardiomyoblast cells induces BMP-dependent hyperacetylation of histone H3 at multiple cardiac gene loci, highlighting a direct link between alcohol exposure and BMP signaling through epigenetic modulation (Shi *et al*., 2017).

In the present study, EtOH exposure at 100 mM for 24 h has shown to induced marked detrimental effects in HL-1 cardiomyocytes. EtOH treatment has significantly enhanced phosphorylation of the BMP pathway transducers SMAD1/5, consistent with earlier observations by Shi et al. (2016), who have reported increased SMAD1/5 phosphorylation in H9c2 cardiomyoblast cells following EtOH exposure (Shi *et al*., 2017). To further evaluate BMP pathway activation, we assessed the expression of key BMP-responsive transcription factors namely *Id1, Gata4, Nkx2.5,* and *Mef2c* which play critical roles at multiple stages of cardiogenesis (Cai *et al*., 2013; Chhabra *et al*., 2019; Qian *et al*., 2014). Our results demonstrated a significant upregulation of these genes at the mRNA level following EtOH treatment, consistent with previous reports (Gao *et al*., 2015; Shi *et al*., 2017). Notably, transcriptome profiling after 100 mM EtOH exposure further manifest the robust overexpression of *Id1* (2.58-fold, *p*=0.0005), *Gata4* (2.0-fold, *p*=0.0366), and *Mef2c* (1.74-fold, *p*=0.0386), as highlighted by the red rectangles in (Figure 6a).

Emerging evidence suggests that BMP2 regulates cardiogenesis-related genes through histone acetyltransferase (HAT) mediated histone acetylation (Zheng *et al*., 2013). In line with this, we have observed a significant increase in HAT activity following EtOH exposure, indicating that EtOH-induced upregulation of BMP signaling is likely mediated, at least in part, through epigenetic histone modifications. To validate the specificity of BMP signaling involvement, the BMP pathway inhibitor LDN-193189, a well-characterized dorsomorphin derivative, is employed. LDN-193189 treatment markedly attenuated EtOH-induced SMAD1/5 phosphorylation, reduced the expression of *Id1*, *Gata4*, *Nkx2.5*, and *Mef2c*, and significantly decreased HAT activity. These findings strongly suggest that LDN-193189 reverses EtOH-mediated BMP pathway activation, thereby confirming a direct role of alcohol in BMP signaling dysregulation.

Consequently, transcriptome profile followed by downstream enrichment analysis reveals 3876 DEGs (≥1.5-fold change, adjusted p<0.05), comprising 295 upregulated and 245 downregulated cardiac-enriched genes in EtOH-treated cells compared to controls group. KEGG pathway enrichment has depicted several upregulated genes are associated with key developmental signaling pathways of gastrulation and body patterning namely TGF-B (Goumans and Ten Dijke, 2018), Wnt (Wiszniak *et al*., 2025), Hippo (Mia and Singh, 2019), Hedgehog, MAPK (Jing-bin *et al*., 2010). Some pathways related to upregulated genes are associated with cellular structure, cell interaction, cell survival such as PI3K-Akt pathway (Ghafouri-Fard *et al*., 2022), ECM-receptor, tight junction and focal adhesion and AGE-RAGE signaling pathway (Wang *et al*., 2025) which is associated with oxidative stress, inflammation, alterations in autophagy flux, and mitochondrial dysfunction. Furthermore, enrichment of adrenergic and calcium signaling pathways suggests disruption of calcium homeostasis and contractile regulation, processes known to increase susceptibility to developmental cardiac abnormalities (Mofid *et al*., 2017). Notably, enhanced TGF-β signaling aligns with our earlier observations of increased SMAD1/5 phosphorylation and upregulation of cardiogenic transcription factors. Similar dysregulations in PI3K-Akt, TGF-β, and Hippo signaling pathways have also been reported in EtOH-induced hiPSC-derived cardiomyocytes, supporting the translational relevance of our observations (Hwang *et al*., 2023). Study by Shepard and Tuma, 2010 have demonstrated that chronic EtOH exposure disrupts the ECM and actin cytoskeleton in both cardiac and non-cardiac cells (Shepard and Tuma, 2010). Such cytoskeletal and ECM alteration has been shown to promote inflammatory responses and impair downstream signaling networks, thereby contributing to broader cellular dysfunction (Suresh and Diaz, 2021).

Conversely, several signaling pathways essential for cardiomyocyte proliferation, differentiation, survival, and immune regulation-including Notch, JAK-STAT (Siddiqui and Mascareno, 2003), MAPK, VEGF (Braile *et al*., 2020), NF-κB (Gutierrez *et al*., 2008), PI3K-Akt, ErbB (Ghafouri-Fard *et al*., 2022), and FoxO signaling (Zhu, 2016) are markedly suppressed. Downregulation of these pathways depicts compromised proliferative potential and perturbed survival signaling, which may adversely affect normal cardiac development. Moreover, reduced activity of pathways involved in maintaining cellular structure and intercellular communication, namely tight junction signaling, highlights disruption of tissue integrity, while dysregulated HIF-1 and p53 signaling reflects hypoxic stress and cellular damage for adaptive responses.

GO analysis further supports these findings by demonstrating that upregulated genes are predominantly associated with heart development, tube morphogenesis, cell differentiation, muscle structure development, ECM organization, sarcolemma formation, and cell-cell junction assembly in biological processes. These genes are also involved in molecular functions such as chromatin binding, protein kinase activity, SMAD and integrin binding. In contrast, downregulated proteins are enriched in negative regulation of apoptosis, circulatory system development, and structural components of the contractile apparatus, including the A-band, I-band, Z-disc, myofibril, and sarcomere. The GO terms related to protein folding chaperone complexes, transcription factor activity, cytoskeletal protein binding, and molecular function regulation indicates impaired proteostasis, contractile integrity, and stress response mechanisms.

Interactome analysis refined with CytoHubba properties has identified key upregulated hub genes, namely *Kras, Fn1, Col1a1, Prkaca, Fbn1, Col6a1, Col6a2, Ccnd1, Col1a2,* and *Myc*, while prominent downregulated hubs genes included *Hsp90aa1, Mdm2, Jun, Hras, Il6, Hsp90ab1, Pdgfra, Cdkn1a, Pparg, Hspa8 and Fos*. In particular, several upregulated hub genes encode collagen components (*Col1a1, Col1a2, Col6a1,* and *Col6a2*), demonstrating extensive ECM remodeling. PAE has been shown to alter collagen subtype composition in the neonatal cardiac ECM (Ninh *et al*., 2019) and pathogenic variants in *COL1A1* are associated with ventricular septal defects and pulmonary atresia (Peters *et al*., 2003). Aberrant COL1A2 expression has been implicated in in right ventricular hypertrophy and heart failure with ToF (Sharma *et al*., 2006). Overexpression of *COL6A1* and *COL6A2* has been linked to atrioventricular septal defects in Down syndrome (Grossman *et al*., 2011; Gramaglia *et al*., 2020). Elevated *Fn1* expression observed in this study is consistent with previous reports of EtOH-induced fibronectin upregulation (CASIN *et al*., 1994). Interestingly, overexpression of this gene has also been observed in right ventricular hypertrophy and failure with ToF (Sharma *et al*., 2006; Arcelli *et al*., 2010) and VSD with PA (Peters *et al*., 2003). *Fbn1*, an essential ECM protein regulating growth factor signaling and tissue architecture, is implicated in Marfan syndrome-associated congenital heart defects (Zodanu *et al*., 2024). The cardiac ECM is an intricately organized scaffold surrounding and interconnecting cardiomyocytes, composed largely of fibrillar collagens that provide mechanical strength, elasticity, and the structural framework required for normal cardiac function. In our study, the coordinated upregulation of major ECM structural components (*Col1a1, Col1a2, Col6a1, Col6a1, Fn1,* and *Fbn1*) suggests excessive ECM deposition and aberrant biomechanical signaling, mechanisms that likely contribute to alcohol-induced congenital cardiac structural defects. K-Ras (*Kras)*, a proto-Oncogene regulate cell growth, differentiation, proliferation, and apoptosis (Bar-Sagi, 2001; Malumbres and Barbacid, 2003). Study by Jason E. Fish, 2020 has revealed that somatic gain-of-function (GoF) KRAS is associated with vascular malformations in mice (Fish *et al*., 2020). Increased cyclin D1-CDK4 (DC) (*Ccnd1)* expression indicates disrupted cell-cycle control and impaired cardiomyocyte proliferation, consistent with observations in dilated cardiomyopathy (Dehghani *et al*., 2023). *Myc* is a master regulatory gene, and its overexpression promotes heightened oxidative stress and aberrant cell-cycle progression, consistent with previous reports (Gu *et al*., 2023). The concurrent upregulation of *Kras, Ccnd1*, and *Myc* points to abnormal cardiomyocyte proliferative signaling and disrupted cell-cycle progression, coupled with elevated oxidative and metabolic stress. Protein Kinase CAMP-Activated Catalytic Subunit Alpha (*Prkaca)* plays a central role in regulating development and metabolism (Palencia-Campos *et al*., 2020) and pathogenic variants have been associated with complex congenital malformations (Sithambaram *et al*., 2024).

Among the downregulated hub genes, *Hsp90aa1* and *Hsp90ab1* encode molecular chaperones essential for protein maturation and cardiac remodeling, and their dysregulation has been associated with myocardial fibrosis and hypertrophy (García *et al*., 2016). *Hspa8* plays a critical role in protein folding and degradation, with altered expression linked to the onset of dilated cardiomyopathy (Chen *et al*., 2016; Portig *et al*., 1997; Liu *et al*., 2025). Decreased molecular chaperones (*Hsp90aa1, Hsp90ab1,* and *Hspa8*) expression further indicates compromised proteostasis and stress adaptation, exacerbating myocardial vulnerability. *HRas*, a member of the Ras family of proto-oncogenes, regulates signaling pathways that drive cardiomyocyte growth and hypertrophic remodeling (Thorburn *et al*., 1993). *Il6* functions as a pleiotropic cytokine produced by both immune and cardiac cells, including endothelial cells, vascular smooth muscle cells, and ischemic cardiomyocytes, plays a crucial role in regulating cardiac metabolism (Kanda and Takahashi, 2004). Interestingly, high IL-6 levels in the serum of children with CHD suggest its potential contribution to pathophysiology of disease (Wang *et al*., 2016; Noori *et al*., 2017). Cardiac defects observed in mutant mouse and zebrafish models support the important role of *Pdgfra* in heart development, (Richarte *et al*., 2007). Notably, *Pdgfra* promotes apoptosis of neural crest cell precursors in both mutant and heterozygous zebrafish models after EtOH exposure and induced developmental abnormalities (McCarthy *et al*., 2013). *Cdkn1a* plays a critical role in cardiac development by integrates inflammatory signaling and apoptotic regulation (Leitch *et al*., 2009). Aberrant activation of *Cdkn1a* driven by *Wdfy3* deficiency disrupts embryonic heart development and results in CHD in mice (Zhang *et al*., 2019). Moreover, loss of *Tbx3* function accelerates cardiomyocyte apoptosis through direct regulation of *Cdkn1a*, further implicating *Cdkn1a* dysregulation in cardiac developmental pathology (Cao *et al*., 2019). *PPARγ* plays a central role in regulating cell growth, migration, and apoptosis while modulating oxidative stress, antioxidant defenses, and inflammatory responses within the cardiovascular system (Chen *et al*., 2008; Oyekan, 2011; Polvani *et al*., 2012). Interestingly, both loss-and gain-of-function approaches including cardiomyocyte-specific and global *PPARγ* knockout as well as agonist treatment have been associated with cardiac hypertrophy and altered blood pressure in mouse models highlighting the necessity of tightly regulated *PPARγ* signaling for cardiac homeostasis. (Duan *et al*., 2005, 2007). Downregulation of developmental and protective signaling nodes (*Pdgfra, Hras, Il6, Cdkn1a,* and *Pparγ*) implicates impaired growth factor-mediated morphogenesis, disrupted inflammatory-metabolic crosstalk, and defective apoptosis regulation during cardiogenesis. *Jun* and *Fos* are proto-oncogenes that heterodimerize to form the AP-1 transcriptional complex in the nucleus, which integrates signals from growth factors, inflammatory mediators, oncogenes, and tumor promoters to regulate cell proliferation (Pfahl and Chytil, 1996). Chronic alcohol exposure has been shown to induce overexpression of AP-1 components (*c-Jun* and *c-Fos*) in vivo (Wang *et al*., 1998). Additionally, suppression of AP-1 transcriptional components (*Jun* and *Fos*) suggests impaired integration of growth, inflammatory, and stress signals during cardiac development. *Mdm2*, a regulator of HIF signaling, contributes to microvascular dysfunction in hypertrophic cardiomyopathy (Shridhar *et al*., 2023). Moreover, qRT-PCR validation of the top hub genes corroborated the network-based findings, further emphasizing their functional relevance in EtOH-induced cardiac toxicity. Collectively, the schematic model integrates our findings into a unified mechanism whereby EtOH exposure aberrantly activates BMP signaling through increased ligand expression, enhanced SMAD1/5 phosphorylation, and HAT-dependent chromatin remodeling. Converging downstream pathways further support a model in which PAE promotes maladaptive ECM remodeling and proliferative imbalance, while simultaneously attenuating key developmental signaling and stress-response mechanisms, ultimately increasing susceptibility to CHD.

Our findings provide clinically relevant mechanistic insight into alcohol exposure-induced congenital cardiac defects by identifying dysregulated BMP signaling components alongside key developmental and metabolic pathways, as well as critical hub genes with potential utility for early risk stratification, biomarker development, and therapeutic intervention aimed at mitigating alcohol-associated cardiotoxicity. Moreover, prior case-control (Burd *et al*., 2007; Martínez-Frías *et al*., 2004; Mateja *et al*., 2012), epidemiological studies (Grewal *et al*., 2008), together with evidence from in-vivo (Serrano *et al*., 2010; Weeks *et al*., 2024), and hiPSCs models (Man *et al*., 2025; Liu *et al*., 2021) further support a causal role for PAE in the development of CHD.

The limitations of this study include the evaluation of EtOH effects exclusively in HL-1 cardiomyocytes, which may not fully recapitulate the complexity of cardiac development in-vivo. In addition, genome-wide chromatin profiling approaches, such as ChIP-seq for histone acetylation or SMAD1/5 occupancy, were not performed to provide direct evidence of epigenetic regulation. Furthermore, the lack of in-vivo validation in animal models or human-relevant system including hiPSC-derived cardiomyocytes or cardiac organoids, limits the translational scope of our findings.

## 5. Conclusion

Our study delineates a comprehensive mechanistic framework linking PAE to disrupted cardiac development. We demonstrate that EtOH (i) aberrantly activates BMP signaling through enhanced SMAD1/5 phosphorylation and HAT-mediated chromatin remodeling, leading to epigenetically driven dysregulated activation of cardiogenic transcriptional programs; (ii) concurrently disrupts multiple developmental, protective, and stress-response pathways that are essential for normal cardiomyocyte maturation and tissue integrity; and (iii) induces pronounced alterations in genes governing ECM remodeling, proliferative balance, and metabolic homeostasis. The coordinated alteration of ECM components, cell-cycle regulators, molecular chaperones, and growth factor-mediated signaling nodes provides a mechanistic basis for perturbed cardiac morphogenesis, compromised proteostasis, and increased susceptibility to oxidative and inflammatory stress. Collectively, these findings establish an integrative molecular model unraveling how PAE predisposes the developing heart to CHD.

## Acknowledgment

We sincerely thank to Prof. Amitabha Bandyopadhyay from Indian Institute of Technology, Kanpur for providing LDN-193189 at the very critical time of experiment. We gratefully acknowledge the University Grants Commission (UGC), Government of India, for awarding Junior and Senior Research Fellowships (577/(CSIR-UGC NET JUNE 2018)) to Jyoti Maddhesiya.

## Data Availability

Raw data and derived data supporting the findings of this study are available from the corresponding author on request.

## Author Contribution

**Bhagyalaxmi Mohapatra**: Conceptualization, Data curation, Formal analysis, Funding acquisition, Project administration, Resources, Supervision, Visualization, Writing – original draft, Writing – review & editing.

**Jyoti Maddhesiya**: Conceptualization, Investigation, Data curation, Formal analysis, Methodology, Software, Validation, Visualization, Writing – original draft, Writing – review & editing.

**Hamim Zafar**: Data curation, Formal analysis, Software, Supervision, Writing-Reviewing and Editing,

**Aditya Gautam**-Data curation, Formal analysis, Methodology, Software, Writing-Reviewing and Editing.

All authors reviewed and approved the final version of the manuscript.

## List of abbreviation

CHD: congenital heart disease
FAS: fetal alcohol syndrome
ASD: atrial septal defect
VSD: ventricular septal defects
NTD: neural tube defects
CLP: cleft lip with or without cleft palate
ToF: tetralogy of Fallot
EAE: embryonic alcohol exposure
hiPSCs: human induced pluripotent stem cells.

