## Supplemental File for "Ethanol-induced activation of BMP signaling and reprogramming of cardiomyocytes transcriptome"

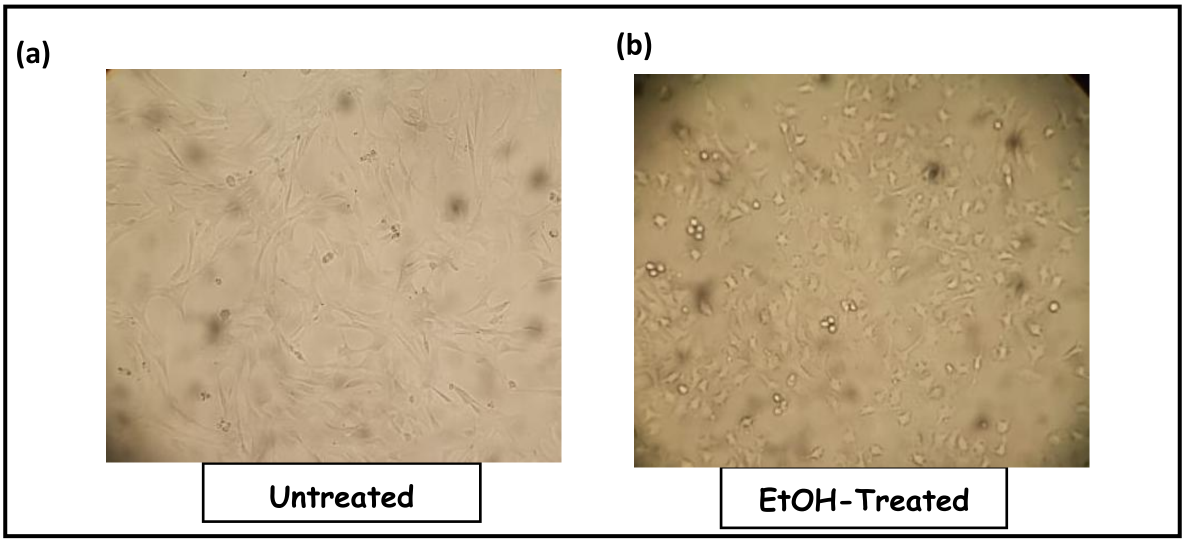


**Supplementary Figure S1** *Phase-contrast microscopy images revealed HL-1 cells* **(a** *Untreated,* **(b)** *EtOH-treated.*
